# Astrocytes in the dorsal vagal complex are not activated by systemic glucoprivation and their chemogenetic activation does not elicit homeostatic glucoregulatory responses in mice

**DOI:** 10.1101/2021.12.15.472746

**Authors:** Alastair J MacDonald, Katherine R Pye, Craig Beall, Kate L J Ellacott

## Abstract

The dorsal vagal complex (DVC) is a brainstem site regulating diverse aspects of physiology including food intake and blood glucose homeostasis. Astrocytes are purported to play an active role in regulating DVC function and, by extension, physiological parameters. Previous work has demonstrated that DVC astrocytes directly sense hormones that regulate food intake and blood glucose and are critical for their effect. In addition, DVC astrocytes in *ex vivo* slices respond to low tissue glucose. The response of neurons, including catecholaminergic neurons, to low glucose is conditional on intact astrocyte signalling in slice preparations suggesting astrocytes are possibly the primary sensors of glucose deprivation (glucoprivation). Based on these findings we hypothesised that if DVC astrocytes act as glucoprivation sensors *in vivo* they would both show a response to systemic glucoprivation and drive physiological responses to restore blood glucose. We found that 2 hours of systemic glucoprivation induced neither FOS nor glial fibrillary acidic protein (GFAP)-immunoreactivity in DVC astrocytes, specifically those in the nucleus of the solitary tract (NTS). Furthermore, we found that while chemogenetic activation of DVC astrocytes suppressed food intake by reducing both meal size and meal number, this manipulation also suppressed food intake under conditions of glucoprivation. Chemogenetic activation of DVC astrocytes did not increase basal blood glucose nor protect against insulin-induced hypoglycaemia. In male mice chemogenetic DVC astrocyte activation did not alter glucose tolerance, in female mice the initial glucose excursion was reduced, suggesting enhanced glucose absorption. Taken together this suggests that as a whole-population DVC astrocytes do not function as glucoprivation sensors *in vivo* in mice. Instead, we propose that DVC astrocytes play an indispensable, homeostatic role to maintain the function of glucoregulatory neuronal circuitry.

## Introduction

The dorsal vagal complex (DVC) is a brain centre involved in regulating many facets of homeostasis including food intake, digestion, cardiovascular reflexes, respiratory reflexes and glucose homeostasis (Finley and Katz, 1992; Machado, 2001; Ritter, Dinh and Li, 2006; Grill and Hayes, 2012; MacDonald and Ellacott, 2020). Situated in the brainstem, the DVC consists of three nuclei: the area postrema (AP; a circumventricular organ), the nucleus of the solitary tract (NTS; a neuronal hub integrating input from the vagus nerves) and the dorsal motor nucleus of the vagus (DMX; containing the cell bodies of preganglionic parasympathetic neurons that comprise the efferent branch of the vagus nerves). Of these constituent nuclei the NTS is the primary sensory integrator of signals from the periphery conveyed by the vagus nerves.

Neurons in the brainstem, particularly those in the NTS and ventrolateral medulla (VLM) are proposed to sense deviations in blood glucose and mediate physiological responses including glucoprivic feeding and hormone secretion to restore blood glucose (Ritter, Dinh and Li, 2006). Local tissue glucoprivation in the NTS and/or VLM induced by injection of 2-deoxyglucose (2-DG; an inhibitor of glycolysis) is sufficient to induce counter-regulatory food intake in rats, suggesting that cells in these sites can directly sense changes in glucose (Ritter, Dinh and Zhang, 2000). Within these brain regions, catecholaminergic neurons, identified by their expression of tyrosine hydroxylase (TH) and/or dopamine beta-hydroxylase (DBH), are a key component of the circuitry underlying glucoprivic responses since their ablation eliminates glucoprivic feeding in rats (Ritter, Bugarith and Dinh, 2001) and their chemogenetic inhibition attenuates glucoprivic feeding in mice (Aklan *et al.*, 2020). In addition, a second population of GABAergic NTS neurons have glucoregulatory capacity since their opto/chemogenetic activation increases blood glucose and glucagon in mice, suggesting the ability of these neurons to drive homeostatic endocrine responses to hypoglycaemia (Lamy *et al.*, 2014; Boychuk *et al.*, 2019).

Although NTS^TH^ neurons are demonstrably required to generate glucoprivic feeding (Ritter, Bugarith and Dinh, 2001; Aklan *et al.*, 2020) it is not clear whether they sense glucose levels cell-autonomously or instead are downstream of other glucose-sensing cells. Indeed, astrocytes in the NTS/DVC have been proposed as the primary cell type underlying low-glucose detection (Rogers and Hermann, 2019). In support of this hypothesis, NTS astrocytes in *ex vivo* brain slices from both rats and mice show increases in intracellular calcium ([Ca^2+^]_i_) in response to low glucose or 2-DG (McDougal *et al.*, 2013; McDougal, Hermann and Rogers, 2013; Rogers *et al.*, 2018). It appears that NTS astrocytes then relay this signal to neighbouring neurons (including NTS^TH^ neurons) by modulating extracellular purine levels (McDougal, Hermann and Rogers, 2013; Rogers *et al.*, 2018). NTS/DVC astrocyte activity and purinergic signalling is required for 2-DG-induced increases in blood glucose in anaesthetised rats, providing functional evidence for astrocyte detection of glucoprivation (Rogers, Ritter and Hermann, 2016). The glucose transporter GLUT2 is proposed to play a glucose-sensing role in astrocytes (Marty *et al.*, 2005; Rogers *et al.*, 2020) as re-expression of glucose transporter GLUT2 in astrocytes of GLUT2^−/−^ mice is sufficient to restore DVC sensitivity to glucoprivation (Marty *et al.*, 2005). In addition, the astrocyte Ca^2+^ response to low glucose in brain slices is abolished with pharmacological GLUT2 blockade (Rogers *et al.*, 2020). Taken together these data suggest NTS/DVC astrocytes detect low glucose, modulate extracellular purine levels (possibly *via* direct release, i.e. gliotransmission) to excite neighbouring neurons and drive physiological responses to restore blood glucose (Rogers and Hermann, 2019).

Thus, evidence from *ex vivo* brain slice experiments and anaesthetised animals suggest astrocytes in the NTS may be the primary sensors of local glucoprivation. Should this be the case, we reasoned that NTS astrocytes would show signs of activation following systemic glucoprivation. Secondly, we hypothesised that the chemogenetic activation of NTS astrocytes, and those in the wider DVC, would drive restorative, physiological responses to glucoprivation for example elevating blood glucose, amplifying glucoprivic feeding and/or attenuating insulin-induced hypoglycaemia.

## Results

### A systemic glucoprivic challenge induced FOS immunoreactivity in the NTS but did not change GFAP-immunoreactive astrocyte number or morphology

In rodent brain slices, NTS astrocytes increase [Ca2^+^]_i_ in response to 2DG-induced glucoprivation (McDougal *et al.*, 2013; McDougal, Hermann and Rogers, 2013; Rogers *et al.*, 2018, 2020) and in anaesthetised rats inhibition of DVC astrocytes suppresses the hyperglycaemic response to systemic 2-DG (Rogers, Ritter and Hermann, 2016). It remains to be seen, however, whether these cells respond to systemic glucoprivation in awake mice.

To assess this question, we injected mice with 2-DG (0.3 g/kg i.p.) and removed food from the cage. 2 hours later mice were perfused (**Figure 1A**). Immunohistochemistry revealed strong induction of the protein product of the immediate early gene FOS in the NTS (**Figure 1B-D**), consistent with previous observations (Ritter, Llewellyn-Smith and Dinh, 1998; Marty *et al.*, 2005). This increase was observed in the NTS proximal to the area postrema with less pronounced differences in the rostral region and no change in caudal sections (**Figure 1D**). Expression of glial fibrillary-acidic protein (GFAP) by NTS astrocytes is dynamic and greater expression and changes in cellular morphology can be indicative of activation (MacDonald and Ellacott, 2020; MacDonald *et al.*, 2020; Escartin *et al.*, 2021). In our experiment, there was no difference between groups in the number of GFAP-immunoreactive cells across the NTS at any of the three rostro-caudal sub-divisions examined (**Figure 1E**). In addition to this there was no difference in morphological complexity, assessed by Sholl analysis, or mean process number of GFAP-immunoreactive cells between groups (**Figure 1F-H**). Finally, FOS was only very rarely co-localised with GFAP-immunoreactive cells and there was no statistically significant difference in the number of co-localised cells between groups (**Figure 1I**). Taken together this suggests that systemic glucoprivation with 2-DG primarily activated neurons in the postremal NTS without altering GFAP-positive astrocyte cytoskeletal protein immunoreactivity, morphology, or immediate early gene immunoreactivity.

**Figure 1.**
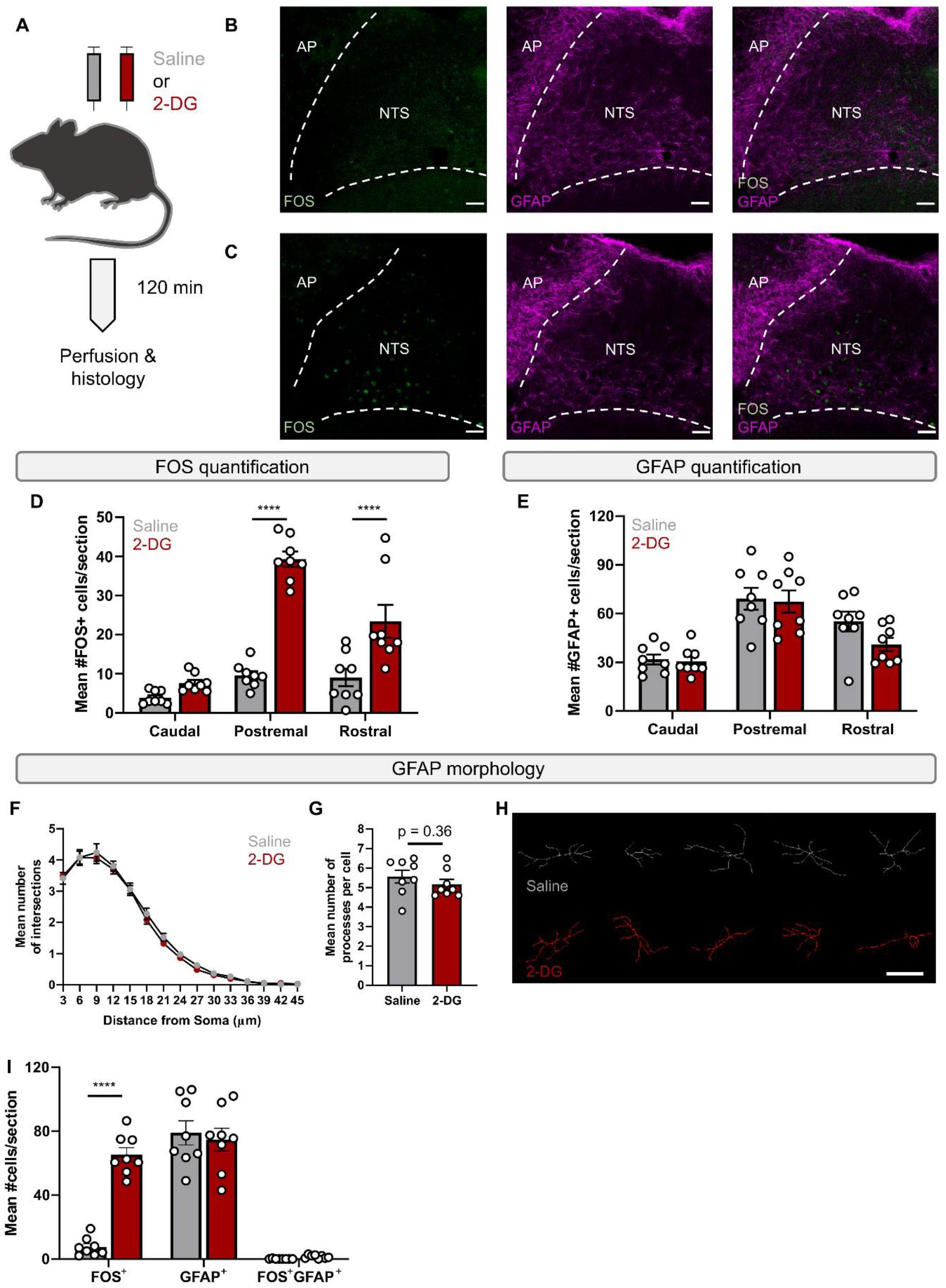
Systemic glucoprivation induced NTS FOS-immunoreactivity but did not change the number of GFAP-immunoreactive cells or their morphology. **A,** Schematic of experimental design (n=8 mice per group, 5 male, 3 female). **B,** Representative image of FOS- and GFAP-immunoreactivity in the NTS of a saline-injected mouse. **C,** Representative image of FOS- and GFAP-immunoreactivity in the NTS of a 2-DG (0.3 g/kg i.p.) injected mouse. **D,** Quantification of the number of FOS cells across the rostro-caudal extent of the NTS (two-way ANOVA with Sidak’s post-hoc test; p_treatment_< 0.0001 F_(1, 42)_= 80.17, p_rostro-caudal position_< 0.0001 F_(2, 42)_= 38.87, p_interaction_<0.0001 F_(2, 42)_= 17.98). **E,** Quantification of the number of GFAP-immunoreactive cells across the rostro-caudal extent of the NTS (two-way ANOVA with Sidak’s post-hoc test; p_treatment_= 0.18 F_(1, 42)_= 1.86, p_rostro-caudal position_< 0.0001 F_(2, 42)_= 25.34, p_interaction_=0.38 F_(2, 42)_= 1.0). **F,** Mean Sholl profile of GFAP-immunoreactive cells in the NTS (n=8 mice per group, 5 male, 3 female. One value calculated per animal as the mean of 10 randomly selected astrocytes. Two-way ANOVA with Sidak’s post-hoc test; p_treatment_= 0.19 F_(1, 210)_= 1.80, p_distance_< 0.0001 F_(14, 210)_= 339.3, p_interaction_=0.99 F_(14, 210)_= 0.25). **G,** Mean number of processes of GFAP-immunoreactive cells in the NTS (n=8 mice per group, 5 male, 3 female. One value calculated per animal as the mean of 10 randomly selected astrocytes. Unpaired t-test). **H,** Representative traces of five cells from a saline- (top row, grey) or 2DG- (bottom row, red) injected mouse. **I,** Quantification of the number of cells immunoreactive for FOS, GFAP or both in the postremal NTS (two-way ANOVA with Sidak’s post-hoc test; p_treatment_< 0.0001 F_(1, 42)_= 23.08, p_target_< 0.0001 F_(2, 42)_= 133.1, p_interaction_<0.0001 F_(2, 42)_= 27.03). Scale bar = 50 μm (**B,C**), 10 μm (**H**). **** = p<0.0001.

These experiments were performed in both male and female mice and the data stratified by sex are shown in **Supplementary figure 1**. In general, effects were consistent across sexes with the exception of FOS in the rostral NTS which was increased in males injected with 2-DG but not females (**Supplementary figure 1B,C**).

### Chemogenetic activation of GFAP-expressing DVC astrocytes suppressed food intake both through reduced meal number and size

We previously demonstrated that chemogenetic activation of GFAP-expressing cells in the DVC with the hM3Dq designer receptor exclusively activated by designer drugs (DREADD) (Armbruster *et al.*, 2007) suppressed food intake in mice (MacDonald *et al.*, 2020). We sought to further characterise this effect by combining this methodology with non-invasive monitoring of food intake, water intake and activity. Mice received bilateral injections of an adeno-associated viral (AAV) vector containing the hM3Dq receptor fused to a fluorescent mCherry reporter under the control of the GFAP promotor (**Figure 2A**). In these mice, mCherry immunoreactivity (IR) was consistently observed in the DVC while the degree of extra-DVC transduction varied between mice (**Figure 2B,C**). At the centre of the injection site (postremal NTS) mCherry-IR was observed in 85.72 ±5.4% of GFAP-IR cells in a diffuse pattern consistent with a membrane-bound protein (**Figure 2D**) (Shigetomi *et al.*, 2013). In the same area, mCherry-IR was absent from NeuN-IR cells (0.00 ±0.00%), indicating the specificity of this approach (**Figure 2E**). Subsequently, we refer to these mice as DVC::GFAP^hM3Dq^ mice.

**Figure 2.**
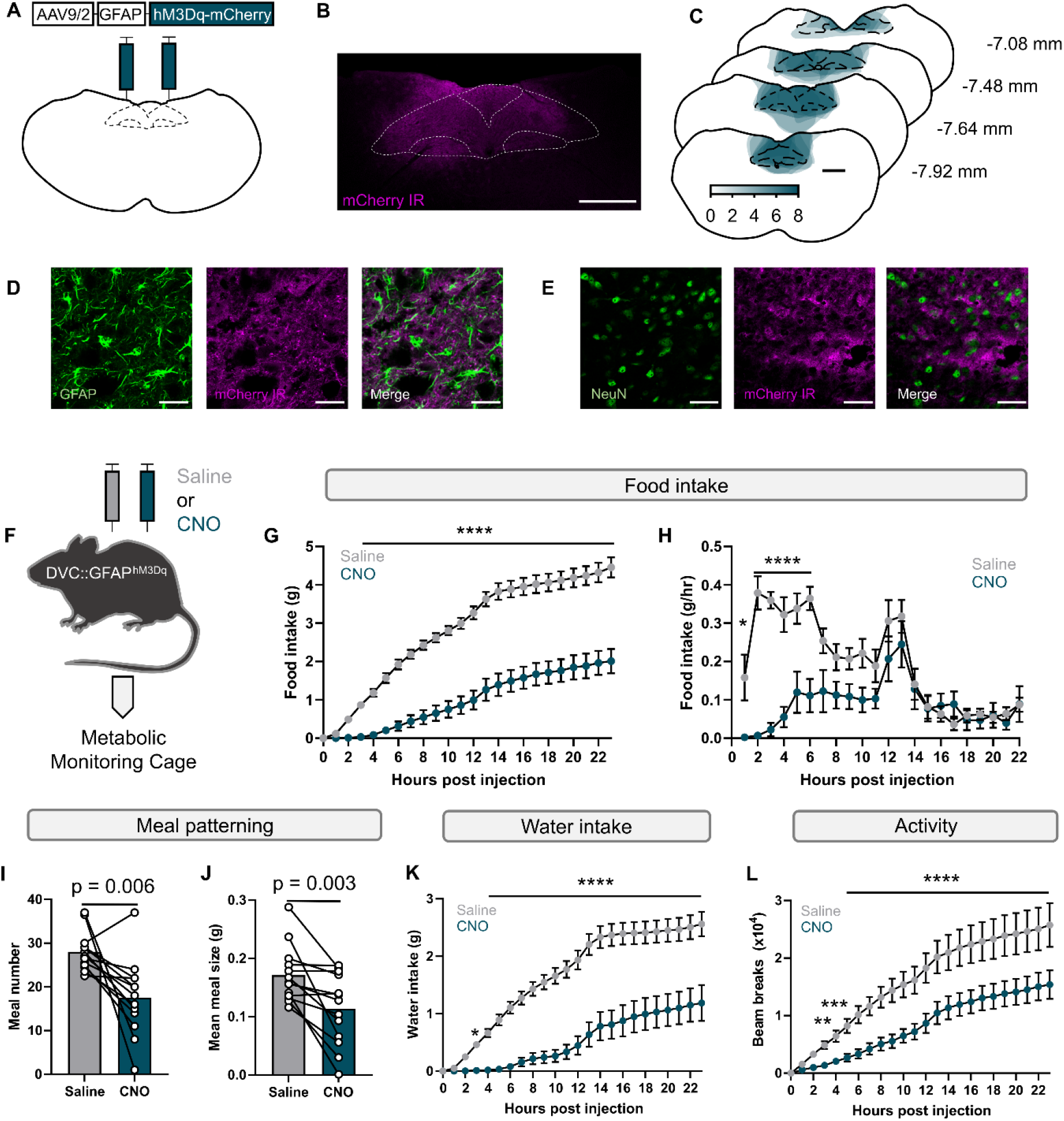
Chemogenetic activation of DVC GFAP-expressing astrocytes reduced food intake by meal number and size. **A,** Schematic of viral vector delivery. **B,** Representative mCherry-immunoreactivity (IR) following viral vector injection. **C,** Quantification of mCherry-IR area (n=8 mice, 5 female, 3 male). **D,** Co-localisation of GFAP- and mCherry-IR in the DVC. **E,** Spatial separation of NeuN and mCherry-IR in the DVC. **F,** Schematic of experimental protocol (n=13 mice, 7 female, 6 male). **G,** Cumulative food intake of DVC::GFAP^hM3Dq^ mice following injection with saline (grey) or CNO (blue) (two-way RM ANOVA with Sidak’s post-hoc test; p_treatment_<0.0001 F_(1, 12)_= 40.39, p_time_< 0.0001 F_(23, 276)_= 184.3, p_interaction_<0.0001 F_(23, 276)_= 20.01). **H,** Food intake rate of DVC::GFAP^hM3Dq^ mice following injection with saline (grey) or CNO (blue) (two-way RM ANOVA with Sidak’s post-hoc test; p_treatment_= 0.0002 F_(1, 12)_= 28.96, ptime< 0.0001 F_(21, 252)_= 7.78, p_interaction_<0.0001 F_(21, 252)_= 7.27). **I,** Meal number of DVC::GFAP^hM3Dq^ mice following injection with saline (grey) or CNO (blue) (paired t-test). **J,** Meal size of DVC::GFAP^hM3Dq^ mice following injection with saline (grey) or CNO (blue) (paired t-test). **K,** Cumulative water intake of DVC::GFAP^hM3Dq^ mice following injection with saline (grey) or CNO (blue) (two-way RM ANOVA with Sidak’s post-hoc test; p_treatment_<0.0001 F_(1, 12)_= 41.10, p_time_< 0.0001 F_(23, 276)_= 68.71, p_interaction_<0.0001 F_(23, 276)_= 13.02). **L**, Cumulative activity of DVC::GFAP^hM3Dq^ mice following injection with saline (grey) or CNO (blue) (two-way RM ANOVA with Sidak’s post-hoc test; p_treatment_<0.0001 F_(1, 12)_= 23.98, p_time_< 0.0001 F_(23, 276)_= 44.18, p_interaction_<0.0001 F_(23, 276)_= 11.45). Scale bar = 500 μm (**B,C**), 25 μm (**D,E**). * = p<0.05, ** = p>0.01, *** = p<0.001, **** = p<0.0001.

DVC::GFAP^hM3Dq^ mice were injected with either saline or CNO (1mg/kg, i.p.) immediately prior to the onset of the dark phase and food intake, water intake and physical activity monitored for the following 23 hours (**Figure 2F**). Injection with CNO reduced cumulative food intake relative to saline injection (**Figure 2G**). Analysis of the rate of food intake shows this effect was most pronounced during the first 6 hours of the dark phase (**Figure 2H**). Analysis of meal patterning revealed that meal number, a metric thought to represent the drive to initiate feeding (i.e., hunger), was reduced following injection with CNO relative to saline (**Figure 2I**). This analysis also showed that the average size of a meal, a metric thought to indicate the strength of meal termination signals (i.e., satiation) was lower following injection with CNO when compared with saline (**Figure 2J**). This suggests that chemogenetic activation of DVC GFAP-expressing astrocytes suppressed food intake both by reducing hunger and advancing satiation.

Water intake was also reduced on CNO injection days as compared with saline (**Figure 2K**). Similarly, activity measured by the cumulative number of beam break events was reduced by injection with CNO relative to saline (**Figure 2L**). As water intake and activity (food seeking) are linked to feeding, from our experiment it is unclear whether these effects are secondary to reduced food intake or driven by distinct mechanisms.

We found that the effects of chemogenetic activation of DVC GFAP-expressing astrocytes was similar in both male and female mice (**Supplementary Figure 2**). In addition, in a separate cohort of mice injected with AAV-GFAP-mCherry, and thus lacking the hM3Dq receptor, treatment with CNO had no effect on food intake, meal patterning or water intake but did induce a slight, but statistically significant, decrease in total activity (**Supplementary Figure 3**).

**Figure 3.**
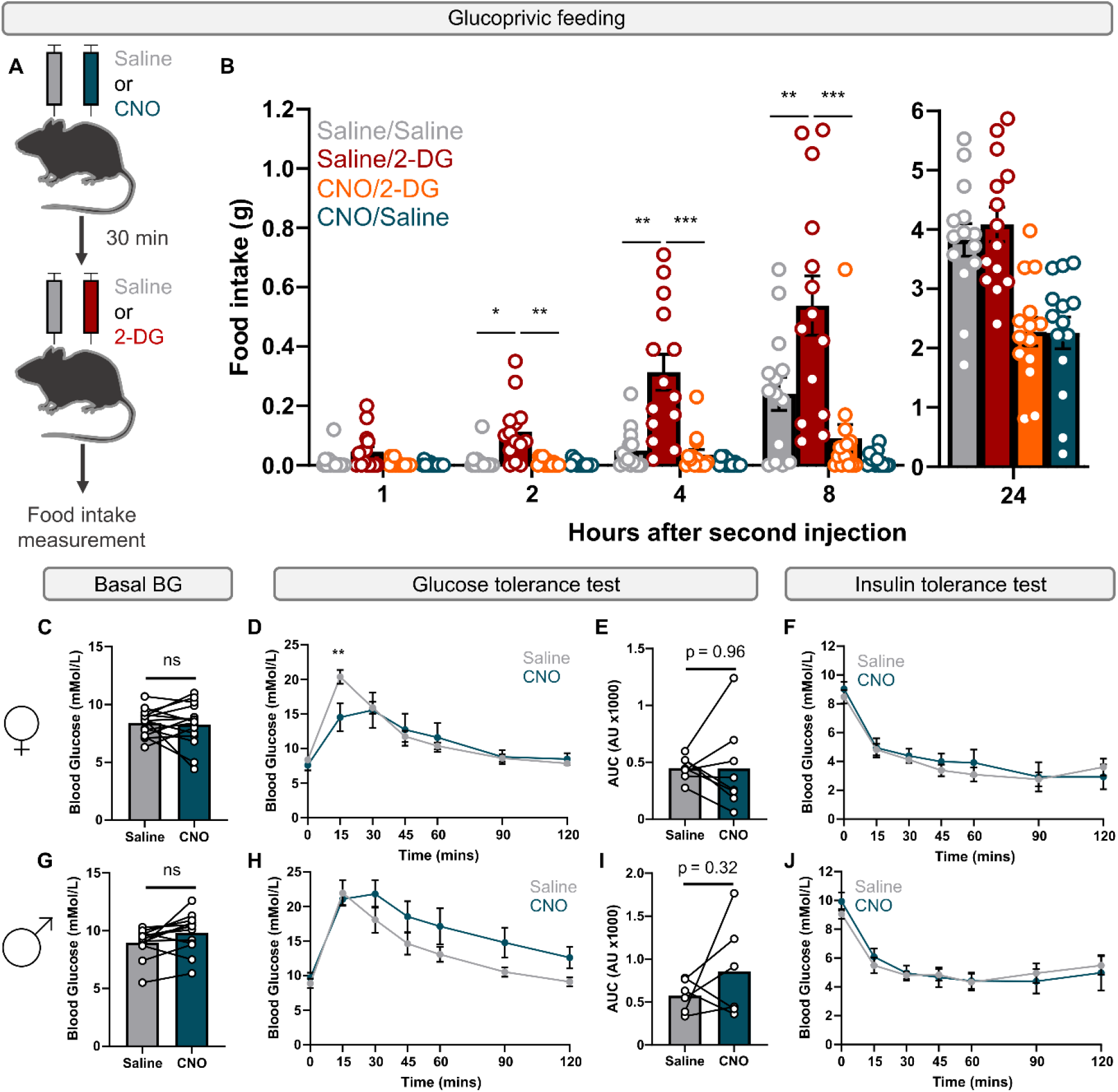
Chemogenetic activation of DVC GFAP-expressing astrocytes attenuated glucoprivic feeding in mice of both sexes and suppressed initial blood glucose excursion in female mice. **A,** Schematic of experimental protocol. **B,** Cumulative food intake of mice injected with Saline or CNO followed by saline or 2-DG (n= 14 mice, 7 female, 7 male, two-way RM ANOVA with Geisser-Greenhouse correction, Sidak’s post-hoc test; p_treatment_< 0.0001 F_(2.672, 34.74)_= 30.40, p_time_< 0.0001 F_(1.079, 14.03)_= 257.9, p_interaction_< 0.0001 F_(2.374, 30.86)_= 16.99). **C,** Basal blood glucose (BG) measured 30 minutes after injection with saline or CNO (n = 15 female mice, paired t-test with Bonferroni correction). **D,** Glucose tolerance curve of mice injected with saline or CNO (n= 8 female mice, two-way RM ANOVA with Sidak’s post-hoc test; p_treatment_= 0.66 F_(1, 7)_= 0.21, p_time_< 0.0001 F_(6, 42)_= 25.37, p_interaction_<0.0001 F_(6, 42)_= 6.24). **E,** Baseline subtracted area under the curve (AUC) for glucose tolerance test in mice injected with saline or CNO (n= 8 female mice, paired t-test). **F,** Insulin tolerance curve of mice injected with saline or CNO (n= 7 female mice, two-way ANOVA with Sidak’s post-hoc test; p_treatment_= 0.69 F_(1, 6)_= 0.18, ptime< 0.0001 F_(6, 36)_= 40.06, p_interaction_=0.59 F_(6, 36)_= 0.78). **G,** Basal blood glucose (BG) measured 30 minutes after injection with saline or CNO (n = 13 male mice, paired t-test with Bonferroni correction). **H,** Glucose tolerance curve of mice injected with saline or CNO (n= 6 male mice, two-way RM ANOVA with Sidak’s post-hoc test; p_treatment_= 0.21 F_(1, 5)_= 2.02, p_time_< 0.0001 F_(6, 30)_= 47.51, p_interaction_=0.21 F_(6, 30)_= 1.52). **I,** Baseline subtracted area under the curve (AUC) for glucose tolerance test in mice injected with saline or CNO (n= 6 male mice, paired t-test). **J,** Insulin tolerance curve of mice injected with saline or CNO (n= 7 male mice, two-way ANOVA with Sidak’s post-hoc test; p_treatment_= 0.89 F_(1, 6)_= 0.02, p_time_< 0.0001 F_(6, 36)_= 24.89, p_interaction_=0.54 F_(6, 36)_= 0.86). * = p<0.05, ** = p>0.01, *** = p<0.001.

### Chemogenetic activation of DVC GFAP-expressing astrocytes suppressed glucoprivic feeding

We reasoned that DVC astrocyte chemogenetic activation may enhance physiological responses to glucoprivation. If so, this could occur by enhancing glucoprivic feeding, elevating basal blood glucose, promoting absorption of glucose from the blood or improving insulin-induced hypoglycaemia. To test the effect of DVC GFAP-expressing astrocyte activation on glucoprivic feeding, DVC::GFAP^hM3Dq^ mice were injected with saline or CNO (1 mg/kg i.p.) during the light phase. 30 minutes later mice were injected with either saline or 2-DG (0.4 g/kg i.p.) and food intake was measured from this time point (**Figure 3A**). Since the effects were consistent across sexes, data were pooled (data are shown separated by sex in **Supplementary figure 4**). In line with other studies, in the absence of chemogenetic activation of DVC GFAP-expressing astrocytes, 2-DG treatment increased food intake compared with injection with saline (Lewis *et al.*, 2006) (**Figure 3B**). In contrast to our hypothesis, injection with CNO prior to 2-DG attenuated glucoprivic feeding to levels equivalent to injection with saline alone (**Figure 3B**). This shows that glucoprivic feeding is suppressed by chemogenetic activation of DVC GFAP-expressing astrocytes.

**Figure 4.**
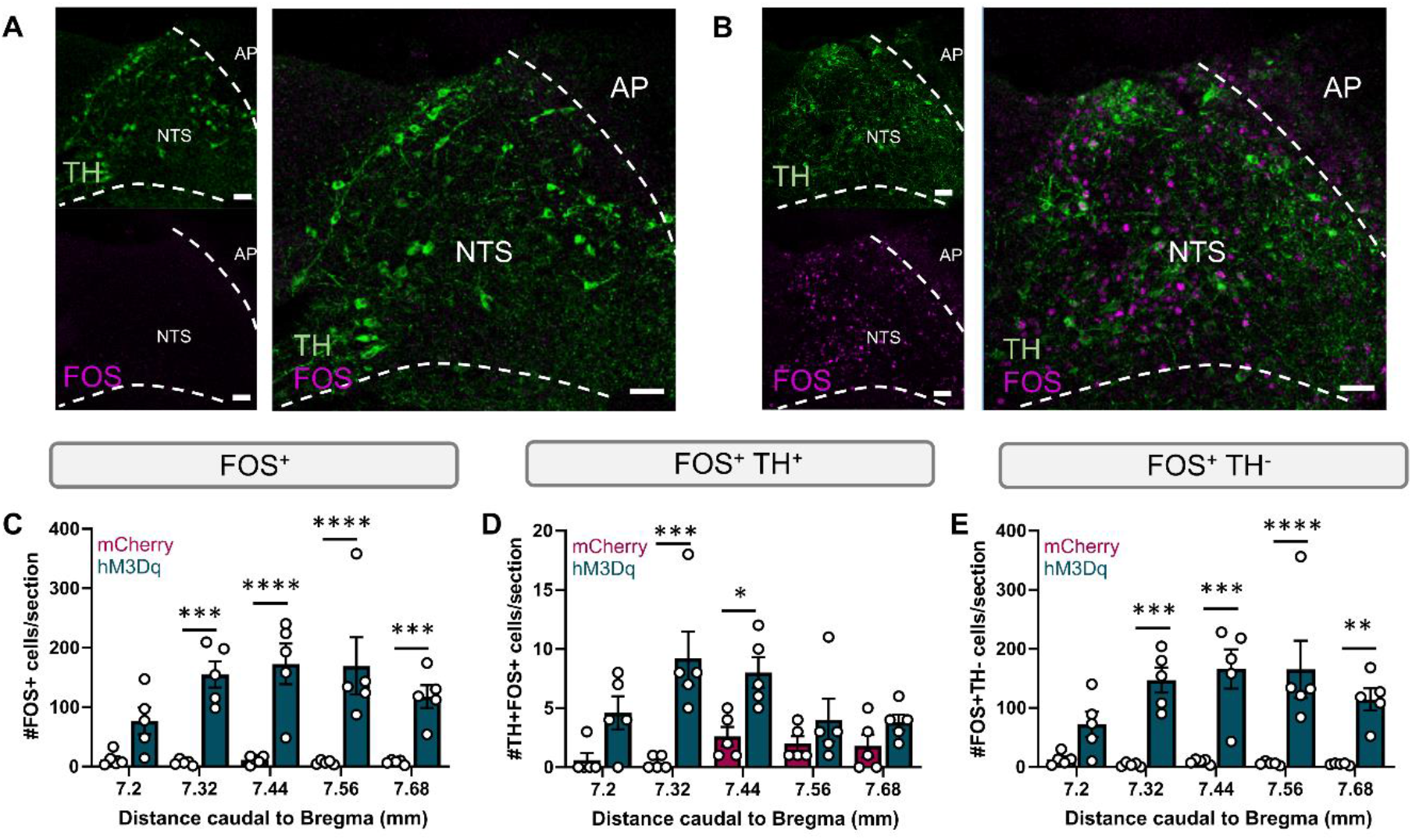
Chemogenetic activation of DVC astrocytes predominantly recruited non-catecholaminergic neurons. **A,** Representative image of TH and FOS-IR in the NTS of DVC::GFAP^mCherry^ mice injected with CNO. **B,** Representative image of TH and FOS-IR in the NTS of DVC::GFAP^hM3Dq^ mice injected with CNO. **C,** Quantification of FOS-IR cells in the NTS of DVC::GFAP^mCherry^ and DVC::GFAP^hM3Dq^ mice injected with CNO (n= 5 mice per group, 2 female, 3 male. Two-way ANOVA with Sidak’s post-hoc test; p_treatment_< 0.0001 F_(1, 40)_= 85.66, p_rostrocaudal position_= 0.19 F_(4, 40)_= 1.60, p_interaction_= 0.15 F_(4, 40)_= 1.81). **D,** Quantification of TH and FOS-IR cells in the NTS of DVC::GFAP^mCherry^ and DVC::GFAP^hM3Dq^ mice injected with CNO (n= 5 mice per group, 2 female, 3 male. Two-way ANOVA with Sidak’s post-hoc test; p_treatment_< 0.0001 F_(1, 40)_= 33.17, prostrocaudal position= 0.10 F_(4, 40)_= 2.11, p_interaction_= 0.04 F_(4, 40)_= 2.69). **E,** Quantification of FOS-IR cells lacking TH-IR in the NTS of DVC::GFAP^mCherry^ and DVC::GFAP^hM3Dq^ mice injected with CNO (n= 5 mice per group, 2 female, 3 male. Two-way ANOVA with Sidak’s post-hoc test; p_treatment_< 0.0001 F_(1, 40)_= 83.20, p_rostrocaudal position_= 0.22 F_(4, 40)_= 1.51, p_interaction_= 0.13 F_(4, 40)_= 1.89). * = p<0.05, ** = p>0.01, *** = p<0.001**** = p<0.0001.

We sought to investigate this further by assessing the effects of chemogenetic activation of DVC GFAP-expressing astrocytes on systemic glucose homeostasis. Activation of GABAergic DVC neurons is sufficient to elevate blood glucose *via* modulation of hepatic glucose production (Boychuk *et al.*, 2019), likely part of the defensive mechanisms against hypoglycaemia. To test whether this mechanism is engaged by chemogenetic activation of DVC astrocytes, blood glucose levels were measured from a cut in the tail 30 minutes after injection with saline or CNO. Neither female nor male DVC::GFAP^hM3Dq^ mice showed a statistically significant difference between treatments (**Figure 3C,G**). Of note, these values are pooled from glucose and insulin tolerance tests (see below) and are therefore corrected for multiple comparisons. These data suggest that, at the time-point tested, activation of DVC GFAP-expressing astrocytes alone was not sufficient to initiate mechanisms to increase blood glucose.

Chemogenetic activation of some DVC neurons (identified by expression of either pre-proglucagon [PPG] or choline acetyltransferase [ChAT]) can improve glucose clearance from the blood following a bolus glucose injection (glucose tolerance) (Shi *et al.*, 2017; NamKoong *et al.*, 2019). We tested whether chemogenetic activation of DVC GFAP-expressing astrocytes was sufficient to recapitulate this effect, potentially *via* these downstream neurons. In female mice, blood glucose was lower 15 minutes following injection with glucose in CNO-injected tests when compared to saline-injected tests (**Figure 3D**). This effect appears to be specific to the initial elevation in blood glucose since there were no statistically significant differences at any other time points (**Figure 3D**) nor in the baseline-subtracted area under the curve (**Figure 3E**). In male mice however, there were no statistically significant effects of CNO on the blood glucose levels at any of the time points (**Figure 3H**), or across the entire test measured by baseline-subtracted area under the curve (**Figure 3i**). This effect in female mice possibly indicates more rapid glucose clearance or enhanced first phase insulin secretion. Due to the divergent effect of CNO between sexes (**Figure 3D,H**) data for all blood glucose measurements were stratified by sex rather than pooled as for other data.

Finally, we tested whether chemogenetic activation of DVC GFAP-expressing astrocytes altered the response to insulin-induced hypoglycaemia. We hypothesised that increasing their activity would be protective against this stimulus by reducing the severity of hypoglycaemia and/or improving blood glucose recovery by stimulating hepatic glucose production (Boychuk *et al.*, 2019). In contrast to our hypothesis, we observed no difference in the response to insulin-induced hypoglycaemia between saline or CNO injection in female or male DVC::GFAP^hM3Dq^ mice neither in the initial reduction in blood glucose nor subsequent recovery (**Figure 3F,J**). Taken together, these results show that activation of DVC GFAP-expressing astrocytes did not modulate basal blood glucose nor was it protective against insulin-induced hypoglycaemia. There was a statistically significant reduction in the peak glucose excursion in female mice during the glucose tolerance test. Finally, activation of DVC GFAP-expressing astrocytes was sufficient to attenuate glucoprivic feeding.

In parallel, we performed these experiments evaluating glucose homeostasis in DVC::GFAP^mCherry^ mice and observed no statistically significant effects of CNO on basal blood glucose, glucose tolerance or insulin tolerance (**Supplementary figure 5 C-J**). We did, however, find a statistically significant increase in food intake in CNO injected mice compared with saline alone 8 hours after injection (**Supplementary figure 5B**). Since we have not observed this previously (**Supplementary figure 3,** MacDonald *et al.*, 2020) it is possible this finding is spurious. Additionally the direction of this finding is opposite to our observation of food intake suppression in DVC::GFAP^hM3Dq^ mice therefore CNO alone does not account for our findings

### Chemogenetic activation of DVC GFAP-expressing astrocytes preferentially activated non-catecholaminergic NTS neurons

We previously showed that chemogenetic activation of DVC GFAP-expressing astrocytes induced FOS immunoreactivity in the DVC and lateral parabrachial nucleus (MacDonald *et al.*, 2020). In the current study we sought to identify whether NTS catecholaminergic neurons, identified by their expression of tyrosine hydroxylase (TH; NTS^TH^ neurons) were among those recruited. NTS^TH^ neurons are involved in both appetite suppression and glucoprivic feeding in mice (Roman, Derkach and Palmiter, 2016; Aklan *et al.*, 2020). Additionally, DVC astrocytes increase [Ca^2+^]_i_ of NTS^TH^ neurons by a purinergic mechanism under low glucose conditions in brain slices (Rogers *et al.*, 2018). DVC::GFAP^hM3Dq^ and DVC::GFAP^mCherry^ mice were injected with CNO and food removed from the cage 2 hours prior to perfusion. Dual-IHC was performed to identify NTS^TH^ neurons and FOS-IR (**Figure 4A,B**). As we found previously, there were far more FOS-IR cells in the NTS of DVC::GFAP^hM3Dq^ mice compared with DVC::GFAP^mCherry^ controls (**Figure 4C**). While there was a modest difference in the number of NTS^TH^ neurons containing FOS-IR between groups (**Figure 4D**), still only a small number of NTS^TH^ neurons had FOS-IR in DVC::GFAP^hM3Dq^ mice relative to the whole NTS^TH^ population. Proportionally, the difference between groups was far greater for FOS-IR cells that did not express TH (**Figure 4E**). This indicates that although there is some recruitment of NTS^TH^ neurons by DVC astrocyte activation this represents only a small fraction of NTS^TH^ neurons and that far more non-TH neurons are recruited.

## Discussion

In this study we set out to determine the ability of GFAP-expressing astrocytes in the NTS and wider DVC to sense systemic glucoprivation and drive responses to restore glucose homeostasis. We found that NTS GFAP-expressing astrocytes do not increase their FOS immunoreactivity following systemic glucoprivation nor do they alter their primary process morphology. Furthermore, the number of GFAP-expressing cells in the NTS was equivalent between groups. In addition, we verified that chemogenetic activation of DVC GFAP-expressing cells suppressed nocturnal food intake in mice of both sexes, and attenuated glucoprivic feeding. We found no effect of chemogenetic DVC GFAP-expressing astrocyte activation on systemic glucose homeostasis save for a modest increase in glucose clearance in female mice. Finally, we observed that chemogenetic activation of DVC GFAP-expressing astrocytes subsequently activates a small number of NTS^TH^ neurons, but these represent a minor proportion of both total NTS^TH^ neurons and total FOS-IR cells suggesting non-TH neurons are predominantly activated.

### NTS astrocytes did not show signs of activation following an acute glucoprivic challenge

In rodent brain slices, NTS astrocytes have been demonstrated to respond directly to both glucoprivation and hormones/hormone receptor ligands that alter systemic glucose homeostasis and food intake (McDougal *et al.*, 2013; McDougal, Hermann and Rogers, 2013; Reiner *et al.*, 2016; Rogers *et al.*, 2018, 2020; Stein *et al.*, 2020). However, in our experiment we did not find any evidence of morphological reorganisation or an increased number of GFAP-immunoreactive cells following 2-DG induced glucoprivation. This does not negate the previous evidence *per se* since astrocytes may show Ca^2+^ fluctuations (resolvable in slice imaging) that do not lead to downstream cellular changes (i.e., GFAP or FOS expression, morphological reorganisation). In the slice experiments only a proportion (~40%) of astrocytes respond to glucoprivation (Rogers *et al.*, 2018) so it is possible that we did not detect an effect if it was ‘diluted’ by non-responsive astrocytes. Both astrocyte metabolism and purinergic transmission are required for hyperglycaemia induced by subcutaneous 2-DG administration in anaesthetised rats (Rogers, Ritter and Hermann, 2016), and while this suggests a primary sensory role for astrocytes, an alternative explanation could be that homeostatic functions of astrocytes, which maintain synaptic transmission in the NTS (MacDonald and Ellacott, 2020), are required for glucose-sensing neurons to mount appropriate physiological responses to rectify deviations from euglycemia.

The time-course we used in our immunohistochemistry experiments (perfusion 2 hours after 2-DG stimulus) is shorter than in our previous studies, which detected a morphological change in NTS astrocytes following 12 hours of high-fat diet intake (MacDonald *et al.*, 2020). So, while it is certainly possible that at a later time point morphological changes may be visible, other studies report rapid reorganisation of astrocyte morphology by food intake or hormone administration on the order of 1-2 hours, albeit using different methodology (electron microscopy) (Nuzzaci *et al.*, 2020; Varela *et al.*, 2021). In order to unify our findings with slice imaging, *in vivo* recording of astrocyte Ca^2+^ with fiber photometry or imaging is a promising future direction (Alhadeff, 2021). Although technical challenges exist, recently fiber photometry recordings have been published for both astrocytes in the striatum and neurons in the NTS (Corkrum *et al.*, 2020; Tan *et al.*, 2020). As such it is possible that combining these approaches may reveal the real-time responses of astrocytes to deviations in energy status in intact animals.

### Chemogenetic activation of DVC GFAP-expressing astrocytes suppressed food intake even under glucoprivic conditions

While chemogenetic activation of DVC GFAP-expressing astrocytes suppresses food intake under conditions of physiological hunger (12 hour overnight fast) (MacDonald *et al.*, 2020), fast-induced food intake and glucoprivic feeding are functionally separable at the circuit level (Hudson and Ritter, 2004). Despite this distinction, chemogenetic DVC GFAP-expressing astrocyte activation suppressed 2-DG induced food intake to levels comparable with saline injection. This suggests that the activation of appetite suppressing neurons downstream of these astrocytes occludes or overrides signals of glucoprivation. Indeed, when we examined the cell types activated by DVC GFAP-expressing astrocyte stimulation only a minority were NTS^TH^ cells, a subpopulation of which drive glucoprivic feeding (Ritter, Dinh and Li, 2006; Aklan *et al.*, 2020). Thus, the net effect of broad chemogenetic activation of DVC GFAP-expressing astrocytes is to suppress food intake. However, this does not rule out the possibility that subsets of DVC astrocytes are specialised for sensing glucoprivation and preferentially communicate with the appropriate neurons to drive responses that restore glucose homeostasis. Further research into the heterogeneity of astrocytes (Batiuk *et al.*, 2020) may reveal markers that allow selective targeting of sub-populations for manipulation and/or monitoring.

### Chemogenetic activation of DVC astrocytes had mild, sex-specific effects on glucose homeostasis

Using chemogenetic activation we uncovered that astrocytes facilitate glucose clearance in female mice. Due to the short onset and duration of this effect it is possible that it is mediated by enhanced first phase glucose-stimulated insulin secretion, although direct measurements of insulin are required to confirm this. This suggests a state-dependence of the effect since, prior to injection of glucose, CNO had no effect on blood glucose compared to saline. Interestingly, this is contrary to predictions made by loss-of-function experiments showing a dependence on DVC astrocytes for 2-DG induced elevations in blood glucose (Rogers, Ritter and Hermann, 2016). Together, our findings and previous work demonstrate that activity of DVC astrocytes is necessary but not sufficient for these elevations in blood glucose. Similarly, the astrocyte-dependent compensatory increase in blood glucose in response to glucoprivation in the anaesthetised rat suggests that activation of astrocytes may be protective against hypoglycaemia (Rogers, Ritter and Hermann, 2016); however, in our study in freely moving mice we found this was not the case, again indicating necessity but not sufficiency of DVC astrocytes for this effect.

We initially posited that DVC astrocytes could be primary glucoprivation sensors, whose activation stimulates counter-regulatory circuitry and downstream responses. However, this divergence of necessity and sufficiency and the absence of overt morphological or immunoreactivity differences in DVC astrocytes following *in vivo* glucoprivation instead suggests that astrocytes in the DVC provide metabolic and/or functional support to counter-regulatory neural circuits that is indispensable to their function.

## Methods

### Mice and housing

C57BL6/J mice were either purchased from Charles River or bred in the University of Exeter Biological Services Unit for experiments. Male and female mice were used for experiments and data were pooled when an effect (or lack thereof) was approximately equal in magnitude and direction in both sexes and/or if control measurements did not differ between sex. Mice were aged > 8 weeks by the time of experiment. Mice were housed in groups of 2-5 with *ad libitum* access to standard laboratory diet (EURodent diet [5LF2], LabDiet) and water in a 12:12 light:dark cycle at 20 ± 2 °C unless otherwise stated.

### Systemic glucoprivation

Mice were individually housed and handled by the experimenter for >6 days prior to the experiment. On the test day mice were injected with either saline or 2DG (0.3 g/kg i.p., Sigma) 2.5 hours into the light phase and food was removed from the cage. Two hours later mice were deeply anaesthetised with sodium pentobarbital and transcardially perfused with saline followed by 4% paraformaldehyde in 0.01M phosphate buffered saline (PBS). Brains were dissected and post-fixed in 4% PFA in 0.01M PBS for 24 hours before being stored in 20% sucrose in 0.01M PBS. Saline or 2DG treatments were randomly assigned and the experimenter was blinded to the group allocations until image analysis was complete. The rostro-caudal ‘bins’ used to quantify cell counts were as follows (all values –mm from Bregma); rostral 6.96 – 7.2, postremal 7.32 – 7.64, caudal 7.76 – 8. Sections were compared to (Paxinos and Franklin, 2008 3^rd^ Edition) to determine the distance from Bregma.

### Viral vector injection surgery

Viral vectors were delivered to the DVC as described previously (Cerritelli *et al.*, 2016; MacDonald *et al.*, 2020). In brief, mice were anaesthetised with ketamine and medetomidine administered as a cocktail. The skin on the back of the head and neck was shaved and the mouse was placed in a stereotaxic frame. Under aseptic conditions, the skin was incised and the muscles parted to expose the atlanato-occipital membrane. This membrane was then incised and a Hamilton needle inserted into the left side of the brain (from obex r/c 0 mm, m/l 0.2mm, d/v 0.5mm) at a 25° angle. 180 nl of viral vector was injected at a rate of 100 nl/min and the needle was kept in place for two minutes after the injection. This process was then repeated on the contralateral (right) side and the wound was closed with sutures in the muscle and skin. The mouse was injected with carprofen and atipamezole (s.c.) and transferred to a heated (24-26°C) recovery cage. The following morning mice received a second injection of carprofen. Mice were given >3 weeks to allow for recovery and viral expression before being used in experiments. Viral vectors used were AAV9/2-hGFAP-hM3Dq-mCherry (ETH Zurich Viral Vector Facility, titre = 5×10^8^ viral genomes/ml) or AAV5/2-hGFAP-mCherry (ViGene Biosciences, titre = 3.89×10^13^) and were diluted 1:3 in sterile saline prior to injection.

### Food intake, water intake and activity monitoring

Mice were individually housed in Promethion metabolic monitoring cages (Sable Systems) which contained three sensors: a mass monitor attached to the food hopper, a mass monitor attached to the water bottle and an XY-beam array surrounding the cage. These permitted the accurate quantification of food intake, water intake and activity using the Promethion system for data acquisition and a macro for data analysis. On the day of the experiment mice were injected with saline or clozapine-*N*-oxide (1 mg/kg, i.p., Tocris) immediately prior to lights off. Data were then acquired for 23 hours. Drug order was not randomised due to the potential duration of CNO-mediated effects and for this reason the experimenter was not blinded. Saline was given first followed by CNO. Following this, the same mice were used for glucose tolerance testing (see below).

### Glucoprivic Feeding

The protocol for assessing glucoprivic feeding was modified from (Lewis *et al.*, 2006). Mice were individually housed and habituated to experimenter handling for > 6 days prior to the experiment. First, mice were injected with saline (i.p.) 3.5 hours into the light phase and received a second injection of saline 30 minutes later. Food intake was then measured manually at 1, 2, 4, 8 and 24 hours later. On the subsequent test days mice were injected with either saline or CNO (1 mg/kg i.p.) 3.5 hours into the light phase followed by 2DG (0.4 g/kg i.p.) 30 minutes later. Food intake was again manually measured at 1, 2, 4, 8 and 24 hours. Test days were separated by at least 72 hours. The allocation of saline or CNO was randomised by coin toss and the investigator was blinded to this allocation until the conclusion of the experiment. On the final day, all mice received CNO (1 mg/kg i.p.) 3.5 hours into the light phase and saline 30 mins later. Food intake was again measured at 1, 2, 4, 8 and 24 hours following the second injection. Following this, the same mice were used for insulin tolerance testing (see below).

### Glucose and Insulin Tolerance Tests

Individually housed mice were transferred to a clean cage with no food 2.5 hours into the light phase. 3.5 hours later the mice were injected with either saline or CNO (1 mg/kg i.p.) and EMLA cream was applied to the tip of the tail. 30 minutes later (6 hours into the light phase) the tip of the tail was cut with scissors and blood glucose measured with a handheld glucometer (Roche). Mice were immediately injected with glucose (2g/kg, Sigma) or insulin (1U/kg, Actrapid NovoNordisk) i.p. and returned to the cage. Blood glucose was measured from the same tail cut 15, 30, 45, 60, 90 and 120 minutes following injection. 10-14 days later the test was repeated with the opposite drug (saline or CNO) given in the first injection. The drug allocation was randomised by coin toss and the investigator was blinded to this allocation until after the experiment concluded.

### Perfusion and histology

All viral-vector injected mice were perfused as described above (“Systemic glucoprivation”) to confirm transduction and quantify viral spread. In some cases, mice were injected with CNO (1 mg/kg i.p.) 2 hours prior to perfusion and food was removed from the cage at this time. Fixed brains were cut into 30 μm thick coronal sections containing the DVC on a freezing sledge microtome (Bright Instruments). For immunohistochemical staining these sections were blocked in 0.01M PBS containing 5% normal donkey serum (Sigma) and 0.3% triton X-100 (Sigma) for 1hr at room temperature, then incubated in a primary antibody (**Table 1**) diluted in 1% normal donkey serum and 0.3% triton in PBS either for 1 hour at room temperature or overnight at 4°C. Sections were then washed 8×5min in PBS and incubated in a secondary antibody (**Table 1**) diluted 1:500 in 0.3% triton in PBS. Sections were then washed 8×5 min in PBS before being mounted onto glass slides and cover-slipped with fluoroshield mounting medium with DAPI (AbCam). Dual staining was performed with sequential primary-secondary incubations. Images were acquired on an upright microscope (Leica AF6000; injection mapping) or a confocal microscope (Leica DMi8; cell counting, co-localisation, morphological analysis) and analysed in FIJI (Schindelin *et al.*, 2012) using simple neurite tracer (SNT) for morphological and Sholl analysis (Ferriera *et al.*, 2014).

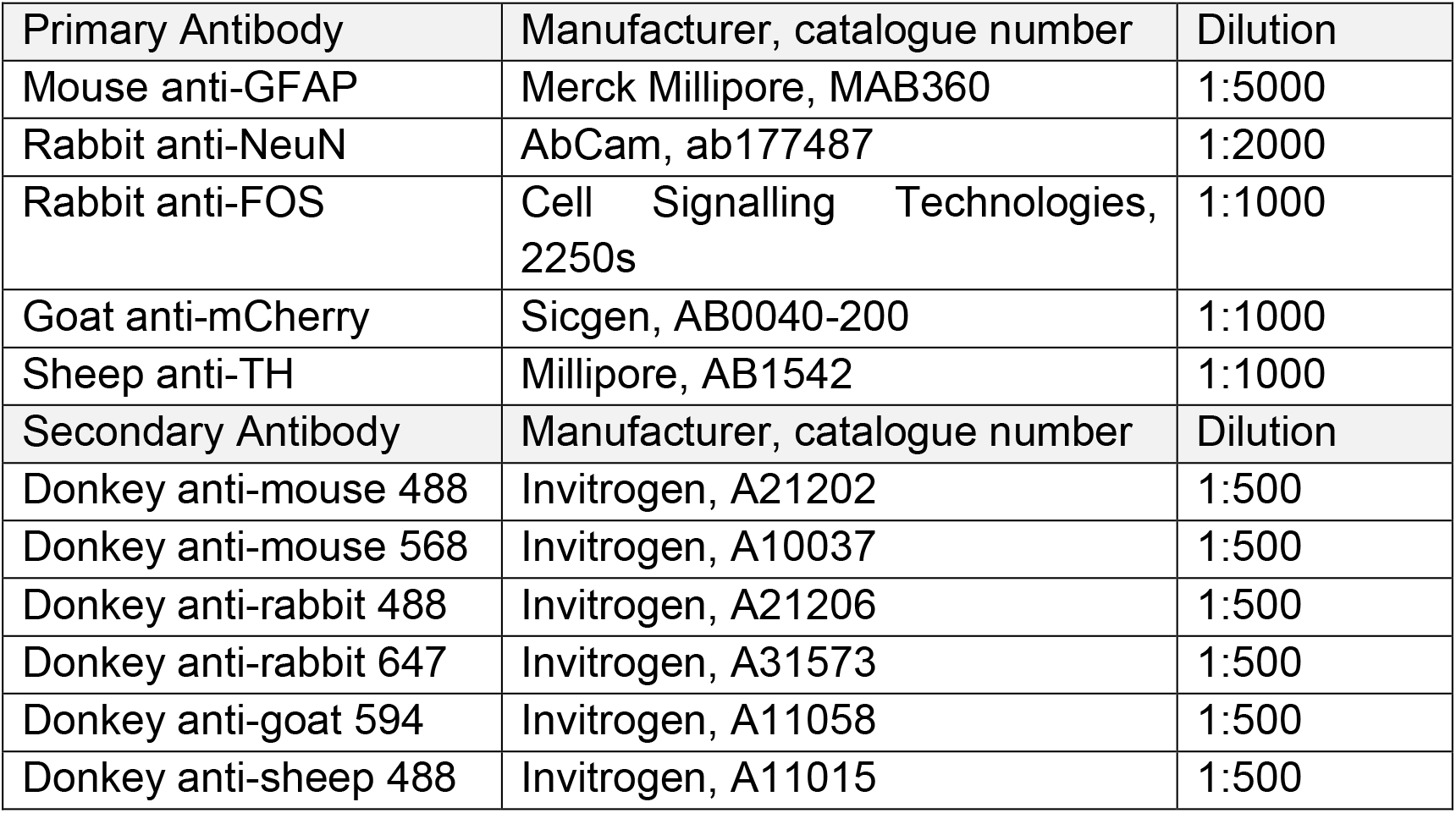

| Primary Antibody | Manufacturer, catalogue number | Dilution |
| --- | --- | --- |
| Mouse anti-GFAP | Merck Millipore, MAB360 | 1:5000 |
| Rabbit anti-NeuN | AbCam, ab177487 | 1:2000 |
| Rabbit anti-FOS | Cell Signalling Technologies, 2250s | 1:1000 |
| Goat anti-mCherry | Sicgen, AB0040-200 | 1:1000 |
| Sheep anti-TH | Millipore, AB1542 | 1:1000 |
| Secondary Antibody | Manufacturer, catalogue number | Dilution |
| Donkey anti-mouse 488 | Invitrogen, A21202 | 1:500 |
| Donkey anti-mouse 568 | Invitrogen, A10037 | 1:500 |
| Donkey anti-rabbit 488 | Invitrogen, A21206 | 1:500 |
| Donkey anti-rabbit 647 | Invitrogen, A31573 | 1:500 |
| Donkey anti-goat 594 | Invitrogen, A11058 | 1:500 |
| Donkey anti-sheep 488 | Invitrogen, A11015 | 1:500 |

### Statistical tests

All values were collated in Prism 9 (GraphPad) for tests of statistical significance (p<0.05). For between-subjects design with one value per subject, unpaired t-tests were used. For within-subjects design with two values per subject, paired t-tests were used. For between-subjects design with multiple values per subject two-way ANOVA was used, with Sidak’s post-hoc test applied to compare between column means. For within-subjects design with two values per subject over time, two-way repeated measure ANOVA was used with Sidak’s post-hoc test applied to compare between column means. For within-subjects design with three or more values per subject over time, two-way repeated measure ANOVA was used with Greenhouse-Geisser correction and Sidak’s post-hoc test applied to compare between column means. No tests for normality were performed and therefore all statistical tests assume a normal distribution. Graphs were generated in Prism, representative images were exported from FIJI and figures were compiled in Inkscape (Inkscape.org). All data are presented as mean ± SEM unless otherwise stated.

## Acknowledgements

The authors would like to thank Matt Isherwood and the staff at the Biological Service Unit at the University of Exeter and Adam Thompson at the RILD building for facilitating this research. The authors would like to thank Ana Miguel Cruz and Paul Weightman Potter for discussions relating to the project. This work was supported by a project grant from Diabetes UK (19/0006035 to KJLE and CB, which funds AJM and KRP). The authors have no conflicts of interest to declare.

## References

Aklan, I. et al. (2020) ‘NTS Catecholamine Neurons Mediate Hypoglycemic Hunger via Medial Hypothalamic Feeding Pathways’, Cell Metabolism, 31(2), pp. 313–326.e5. doi: 10.1016/j.cmet.2019.11.016.

Alhadeff, A. L. (2021) ‘Monitoring in Vivo Neural Activity to Understand Gut-Brain Signaling’, Endocrinology (United States), 162(5), pp. 1–12. doi: 10.1210/endocr/bqab029.

Armbruster, B. N. et al. (2007) ‘Evolving the lock to fit the key to create a family of G protein-coupled receptors potently activated by an inert ligand’, Proceedings of the National Academy of Sciences of the United States of America, 104(12), pp. 5163–5168. doi: 10.1073/pnas.0700293104.

Batiuk, M. Y. et al. (2020) ‘Identification of region-specific astrocyte subtypes at single cell resolution’, Nature Communications, 11(1), pp. 1–15. doi: 10.1038/s41467-019-14198-8.

Boychuk, C. R. et al. (2019) ‘A hindbrain inhibitory microcircuit mediates vagally-coordinated glucose regulation’, Scientific Reports, 9(1), pp. 1–12. doi: 10.1038/s41598-019-39490-x.

Cerritelli, S. et al. (2016) ‘Activation of brainstem proopiomelanocortin neurons produces opioidergic analgesia, bradycardia and bradypnoea’, PLoS ONE, 11(4), pp. 1–26. doi: 10.1371/journal.pone.0153187.

Corkrum, M. et al. (2020) ‘Dopamine-Evoked Synaptic Regulation in the Nucleus Accumbens Requires Astrocyte Activity’, Neuron, 105(6), pp. 1036–1047.e5. doi: 10.1016/j.neuron.2019.12.026.

Escartin, C. et al. (2021) ‘Reactive astrocyte nomenclature, definitions, and future directions’, Nature Neuroscience, 24(3), pp. 312–325. doi: 10.1038/s41593-020-00783-4.

Ferriera, T. A. et al. (2014) ‘Neuronal morphometry directly from bitmap images’, Nature Methods, 11(10), pp. 981–984. doi: 10.1038/nmeth.3102.

Finley, J. C. W. and Katz, D. M. (1992) ‘The central organization of carotid body afferent projections to the brainstem of the rat’, Brain Research, 572(1–2), pp. 108–116. doi: 10.1016/0006-8993(92)90458-L.

Grill, H. J. and Hayes, M. R. (2012) ‘Hindbrain neurons as an essential hub in the neuroanatomically distributed control of energy balance’, Cell Metabolism, 16(3), pp. 296–309. doi: 10.1016/j.cmet.2012.06.015.

Hudson, B. and Ritter, S. (2004) ‘Hindbrain catecholamine neurons mediate consummatory responses to glucoprivation’, Physiology and Behavior, 82(2–3), pp. 241–250. doi: 10.1016/j.physbeh.2004.03.032.

Lamy, C. M. et al. (2014) ‘Hypoglycemia-activated GLUT2 neurons of the nucleus tractus solitarius stimulate vagal activity and glucagon secretion’, Cell Metabolism, 19(3), pp. 527–538. doi: 10.1016/j.cmet.2014.02.003.

Lewis, S. R. et al. (2006) ‘Genetic variance contributes to ingestive processes: A survey of 2-deoxy-D-glucose-induced feeding in eleven inbred mouse strains’, Physiology and Behavior, 87(3), pp. 595–601. doi: 10.1016/j.physbeh.2005.12.002.

MacDonald, A. J. et al. (2020) ‘Regulation of food intake by astrocytes in the brainstem dorsal vagal complex’, Glia, 68(6), pp. 1241–1254. doi: 10.1002/glia.23774.

MacDonald, A. J. and Ellacott, K. L. J. (2020) ‘Astrocytes in the nucleus of the solitary tract: Contributions to neural circuits controlling physiology’, Physiology and Behavior, 223(November 2019), p. 112982. doi: 10.1016/j.physbeh.2020.112982.

Machado, B. H. (2001) ‘Neurotransmission of the cardiovascular reflexes in the nucleus tractus solitarii of awake rats’, Annals of the New York Academy of Sciences, 940, pp. 179–196. doi: 10.1111/j.1749-6632.2001.tb03676.x.

Marty, N. et al. (2005) ‘Regulation of glucagon secretion by glucose transporter type 2 (glut2) and astrocyte dependent glucose sensors’, Journal of Clinical Investigation, 115(12), pp. 3545–3553. doi: 10.1172/JCI26309.

McDougal, D. H. et al. (2013) ‘Astrocytes in the hindbrain detect glucoprivation and regulate gastric motility’, Autonomic Neuroscience: Basic and Clinical, 175(1–2), pp. 61–69. doi: 10.1016/j.autneu.2012.12.006.

McDougal, D. H., Hermann, G. E. and Rogers, R. C. (2013) ‘Astrocytes in the nucleus of the solitary tract are activated by low glucose or glucoprivation: Evidence for glial involvement in glucose homeostasis’, Frontiers in Neuroscience, 7(7 DEC), pp. 1–10. doi: 10.3389/fnins.2013.00249.

NamKoong, C. et al. (2019) ‘Chemogenetic manipulation of parasympathetic neurons (DMV) regulates feeding behavior and energy metabolism’, Neuroscience Letters, 712(June), p. 134356. doi: 10.1016/j.neulet.2019.134356.

Nuzzaci, D. et al. (2020) ‘Postprandial Hyperglycemia Stimulates Neuroglial Plasticity in Hypothalamic POMC Neurons after a Balanced Meal’, Cell Reports, 30(9), pp. 3067–3078.e5. doi: 10.1016/j.celrep.2020.02.029.

Reiner, D. J. et al. (2016) ‘Astrocytes regulate GLP-1 receptor-mediated effects on energy balance’, Journal of Neuroscience, 36(12), pp. 3531–3540. doi: 10.1523/JNEUROSCI.3579-15.2016.

Ritter, S., Bugarith, K. and Dinh, T. T. (2001) ‘Immunotoxic destruction of distinct catecholamine subgroups produces selective impairment of glucoregulatory responses and neuronal activation’, Journal of Comparative Neurology, 432(2), pp. 197–216. doi: 10.1002/cne.1097.

Ritter, S., Dinh, T. T. and Li, A. J. (2006) ‘Hindbrain catecholamine neurons control multiple glucoregulatory responses’, Physiology and Behavior, 89(4), pp. 490–500. doi: 10.1016/j.physbeh.2006.05.036.

Ritter, S., Dinh, T. T. and Zhang, Y. (2000) ‘Localization of hindbrain glucoreceptive sites controlling food intake and blood glucose’, Brain Research, 856(1–2), pp. 37–47. doi: 10.1016/S0006-8993(99)02327-6.

Ritter, S., Llewellyn-Smith, I. and Dinh, T. T. (1998) ‘Subgroups of hindbrain catecholamine neurons are selectively activated by 2-deoxy-D-glucose induced metabolic challenge’, Brain Research, 805(1–2), pp. 41–54. doi: 10.1016/S0006-8993(98)00655-6.

Rogers, R. C. et al. (2018) ‘Response of catecholaminergic neurons in the mouse hindbrain to glucoprivic stimuli is astrocyte dependent’, American Journal of Physiology - Regulatory Integrative and Comparative Physiology, 315(1), pp. R153–R164. doi: 10.1152/ajpregu.00368.2017.

Rogers, R. C. et al. (2020) ‘Evidence that hindbrain astrocytes in the rat detect low glucose with a glucose transporter 2-phospholipase C-calcium release mechanism’, American Journal of Physiology - Regulatory Integrative and Comparative Physiology, 318(1), pp. R38–R48. doi: 10.1152/AJPREGU.00133.2019.

Rogers, R. C. and Hermann, G. E. (2019) ‘Hindbrain astrocytes and glucose counter-regulation’, Physiology and Behavior, 204(February), pp. 140–150. doi: 10.1016/j.physbeh.2019.02.025.

Rogers, R. C., Ritter, S. and Hermann, G. E. (2016) ‘Hindbrain cytoglucopenia-induced increases in systemic blood glucose levels by 2-deoxyglucose depend on intact astrocytes and adenosine release’, American Journal of Physiology - Regulatory Integrative and Comparative Physiology, 310(11), pp. R1102–R1108. doi: 10.1152/ajpregu.00493.2015.

Roman, C. W., Derkach, V. A. and Palmiter, R. D. (2016) ‘Genetically and functionally defined NTS to PBN brain circuits mediating anorexia’, Nature Communications, 7(May), pp. 1–11. doi: 10.1038/ncomms11905.

Schindelin, J. et al. (2012) ‘Fiji: An open-source platform for biological-image analysis’, Nature Methods, 9(7), pp. 676–682. doi: 10.1038/nmeth.2019.

Shi, X. et al. (2017) ‘Acute activation of GLP-1-expressing neurons promotes glucose homeostasis and insulin sensitivity’, Molecular Metabolism, 6(11), pp. 1350–1359. doi: 10.1016/j.molmet.2017.08.009.

Shigetomi, E. et al. (2013) ‘Imaging calcium microdomains within entire astrocyte territories and endfeet with GCaMPs expressed using adeno-associated viruses’, Journal of General Physiology, 141(5), pp. 633–647. doi: 10.1085/jgp.201210949.

Stein, L. M. et al. (2020) ‘Dorsal vagal complex and hypothalamic glia differentially respond to leptin and energy balance dysregulation’, Translational Psychiatry, 10(1). doi: 10.1038/s41398-020-0767-0.

Tan, H. E. et al. (2020) ‘The gut–brain axis mediates sugar preference’, Nature, 580(7804), pp. 511–516. doi: 10.1038/s41586-020-2199-7.

Varela, L. et al. (2021) ‘Hunger-promoting AgRP neurons trigger an astrocyte-mediated feed-forward autoactivation loop in mice’, Journal of Clinical Investigation, 131(10). doi: 10.1172/JCI144239.

