## supplementary for "Astrocytes in the dorsal vagal complex are not activated by systemic glucoprivation and their chemogenetic activation does not elicit homeostatic glucoregulatory responses in mice"

### Supplementary Material

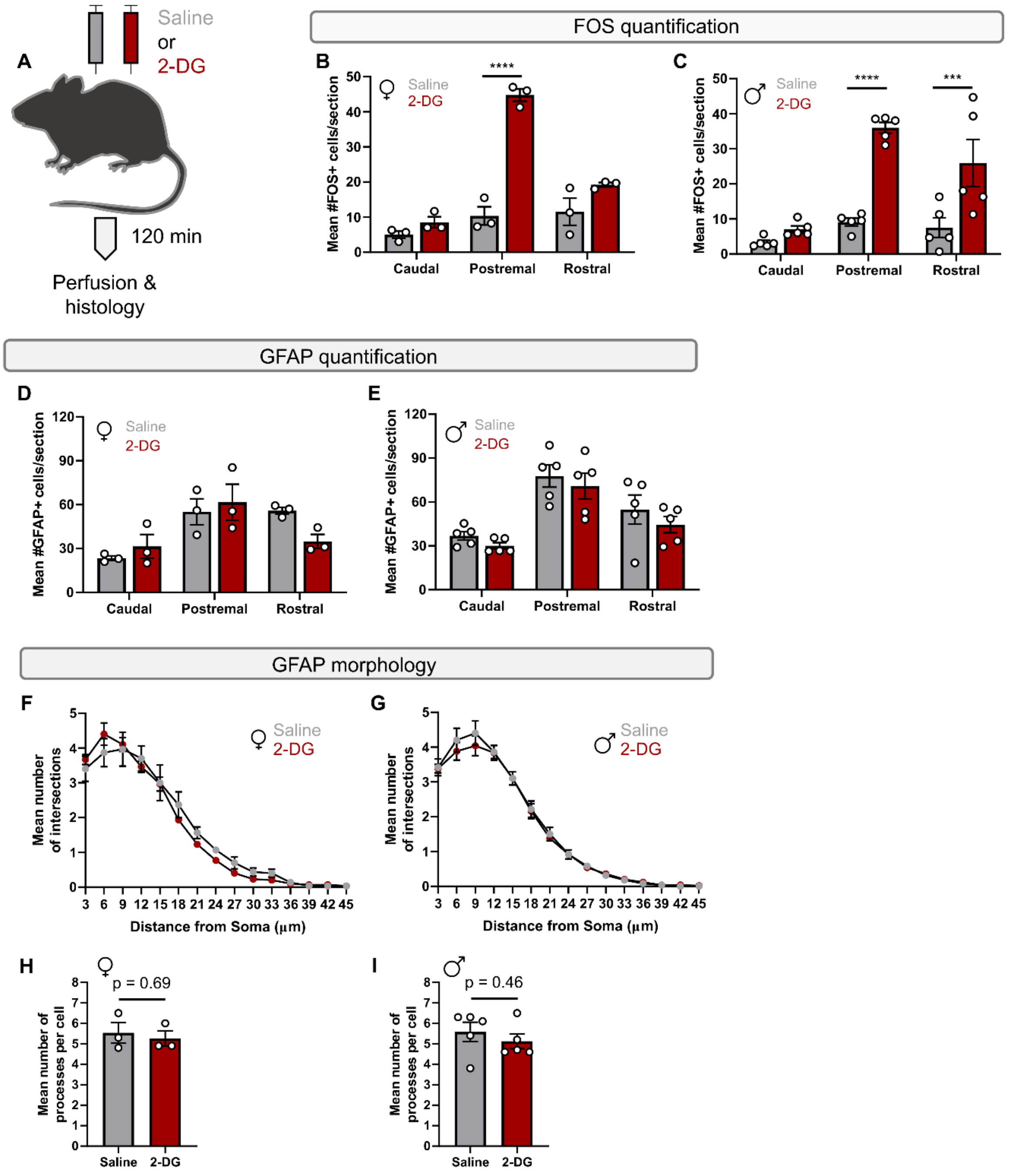


**Supplementary Figure 1 | Systemic glucoprivation effects on FOS-immunoreactivity, GFAP-immunoreactivity and GFAP-immunoreactive cell morphology stratified by sex. A,** Schematic of experimental design (n=3 mice per group [females], n=5 mice per group [males]). **B,** Quantification of the number of FOS-immunoreactive cells across the rostro-caudal extent of the NTS in female mice (two-way ANOVA with Sidak's post-hoc test; $p_{treatment} < 0.0001$ $F_{(1,\ 12)} = 73.37$, $p_{rostrocaudal\ position} < 0.0001$ $F_{(2,\ 12)} = 46.13$, $p_{interaction} < 0.0001$ $F_{(2,\ 12)} = 29.68$). **C,** Quantification of the number of

FOS-immunoreactive cells across the rostro-caudal extent of the NTS in male mice (two-way ANOVA with Sidak's post-hoc test; $p_{treatment} < 0.0001$ $F_{(1, 24)} = 41.80$, $p_{rostrocaudal\ position} < 0.0001$ $F_{(2, 24)} = 16.31$, $p_{interaction} = 0.004$ $F_{(2, 24)} = 7.09$). **D,** Quantification of the number of GFAP cells across the rostro-caudal extent of the NTS in female mice (two-way ANOVA with Sidak's post-hoc test; $p_{treatment} = 0.73$ $F_{(1, 12)} = 0.13$, $p_{rostrocaudal\ position} = 0.004$ $F_{(2, 12)} = 8.88$, $p_{interaction} = 0.13$ $F_{(2, 12)} = 2.45$). **E,** Quantification of the number of GFAP cells across the rostro-caudal extent of the NTS in male mice (two-way ANOVA with Sidak's post-hoc test; $p_{treatment} = 0.16$ $F_{(1, 24)} = 2.07$, $p_{rostrocaudal\ position} < 0.0001$ $F_{(2, 24)} = 18.20$, $p_{interaction} = 0.96$ $F_{(2, 24)} = 0.04$). **F,** Mean Sholl profile of GFAP-immunoreactive cells in the NTS of female mice (One value calculated per animal as the mean of 10 randomly selected astrocytes. Two-way ANOVA with Sidak's post-hoc test; $p_{treatment} = 0.37$ $F_{(1, 60)} = 0.8$, $p_{distance} < 0.0001$ $F_{(14, 60)} = 110.3$, $p_{interaction} = 0.78$ $F_{(14, 60)} = 0.69$). **G,** Mean Sholl profile of GFAP-immunoreactive cells in the NTS of male mice (One value calculated per animal as the mean of 10 randomly selected astrocytes. Two-way ANOVA with Sidak's post-hoc test; $p_{treatment} = 0.11$ $F_{(1, 120)} = 2.61$, $p_{distance} < 0.0001$ $F_{(14, 120)} = 173.2$, $p_{interaction} = 0.99$ $F_{(14, 120)} = 0.24$). **H**, Mean number of processes of GFAP-immunoreactive cells in the NTS of female mice (One value calculated per animal as the mean of 10 randomly selected astrocytes. Unpaired t-test). **I**, Mean number of processes of GFAP-immunoreactive cells in the NTS of male mice (One value calculated per animal as the mean of 10 randomly selected astrocytes. Unpaired t-test).

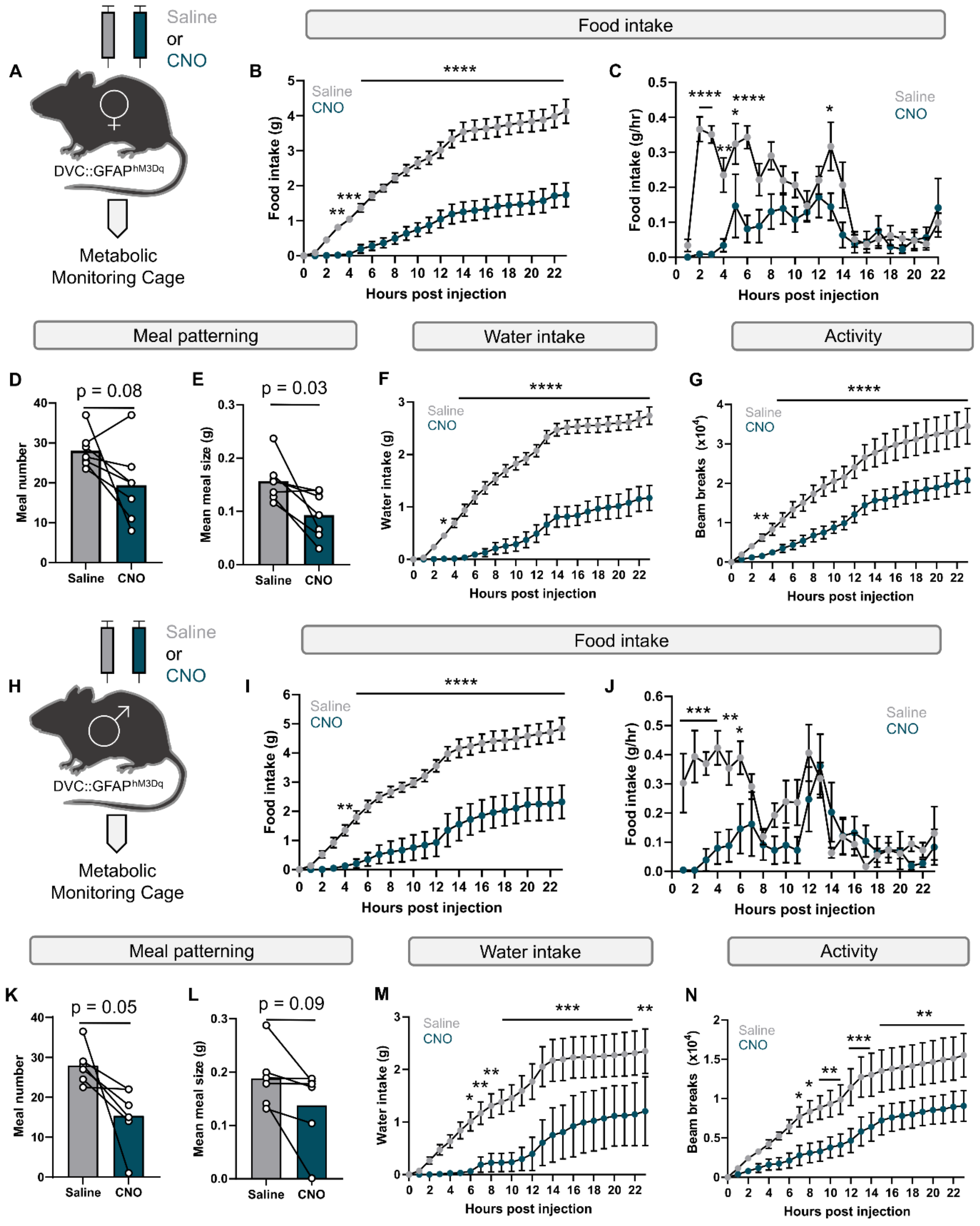


**Supplementary figure 2 | Chemogenetic activation of DVC GFAP-expressing astrocytes: effects on food intake, meal patterning, water intake and activity stratified by sex. A,** Schematic of experimental design (n= 7 female mice). **B,** Cumulative food intake of female DVC::GFAP$^{hM3Dq}$ mice following injection with saline (grey) or CNO (blue) (two-way RM ANOVA with Sidak's post-hoc test; $p_{treatment}$= 0.0025 $F_{(1, 6)}$= 40.39, $p_{time}$< 0.0001 $F_{(23, 138)}$= 154.1, $p_{interaction}$<0.0001 $F_{(23, 138)}$= 11.95). **C,** Food intake rate of female DVC::GFAP$^{hM3Dq}$ mice following injection with saline (grey) or CNO (blue) (two-way RM ANOVA with Sidak's post-hoc test; $p_{treatment}$<0.007 $F_{(1, 6)}$= 16.11, $p_{time}$< 0.0001 $F_{(21, 126)}$= 7.13, $p_{interaction}$<0.0001 $F_{(21, 126)}$= 4.71). **D,** Meal number of female DVC::GFAP$^{hM3Dq}$ mice following injection with saline (grey) or CNO (blue) (paired t-test). **E,** Meal size of female DVC::GFAP$^{hM3Dq}$ mice following injection with saline (grey) or CNO (blue) (paired t-test). **F,** Cumulative water intake of female DVC::GFAP$^{hM3Dq}$ mice following injection with saline (grey) or CNO (blue) (two-way RM ANOVA with Sidak's post-hoc test; $p_{treatment}$= 0.0002 $F_{(1, 6)}$= 61.97, $p_{time}$< 0.0001 $F_{(23, 138)}$= 121.8, $p_{interaction}$<0.0001 $F_{(23, 138)}$= 20.39). **G,** Cumulative activity of male DVC::GFAP$^{hM3Dq}$ mice following injection with saline (grey) or CNO (blue) (two-way RM ANOVA with Sidak's post-hoc test; $p_{treatment}$= 0.001 $F_{(1, 6)}$= 35.47, $p_{time}$< 0.0001 $F_{(23, 138)}$= 51.48, $p_{interaction}$<0.0001 $F_{(23, 138)}$= 13.89). **H,** Schematic of experimental design (n=6 male mice). **I,** Cumulative food intake of male DVC::GFAP$^{hM3Dq}$ mice following injection with saline (grey) or CNO (blue) (two-way RM ANOVA with Sidak's post-hoc test; $p_{treatment}$= 0.012 $F_{(1, 5)}$= 14.88, $p_{time}$< 0.0001 $F_{(23, 115)}$= 81.49, $p_{interaction}$<0.0001 $F_{(23, 115)}$= 7.59). **J,** Food intake rate of male DVC::GFAP$^{hM3Dq}$ mice following injection with saline (grey) or CNO (blue) (two-way RM ANOVA with Sidak's post-hoc test; $p_{treatment}$= 0.02 $F_{(1, 5)}$= 11.09, $p_{time}$< 0.0001 $F_{(21, 105)}$= 3.57, $p_{interaction}$<0.0001 $F_{(21, 105)}$= 4.97). **K,** Meal number of male DVC::GFAP$^{hM3Dq}$ mice following injection with saline (grey) or CNO (blue) (paired t-test). **L,** Meal size of male DVC::GFAP$^{hM3Dq}$ mice following injection with saline (grey) or CNO (blue) (paired t-test). **M,** Cumulative water intake of male DVC::GFAP$^{hM3Dq}$ mice following injection with saline (grey) or CNO (blue) (two-way RM ANOVA with Sidak's post-hoc test; $p_{treatment}$= 0.03 $F_{(1, 5)}$= 8.32, $p_{time}$< 0.0001 $F_{(23, 115)}$= 14.73, $p_{interaction}$=0.001 $F_{(23, 115)}$= 2.43). **N,** Cumulative activity of male DVC::GFAP$^{hM3Dq}$ mice following injection with saline (grey) or CNO (blue) (two-way RM ANOVA with Sidak's post-hoc test; $p_{treatment}$= 0.1 $F_{(1, 5)}$= 4.05, $p_{time}$< 0.0001 $F_{(23, 115)}$= 65.07, $p_{interaction}$=0.02 $F_{(23, 115)}$= 1.80).

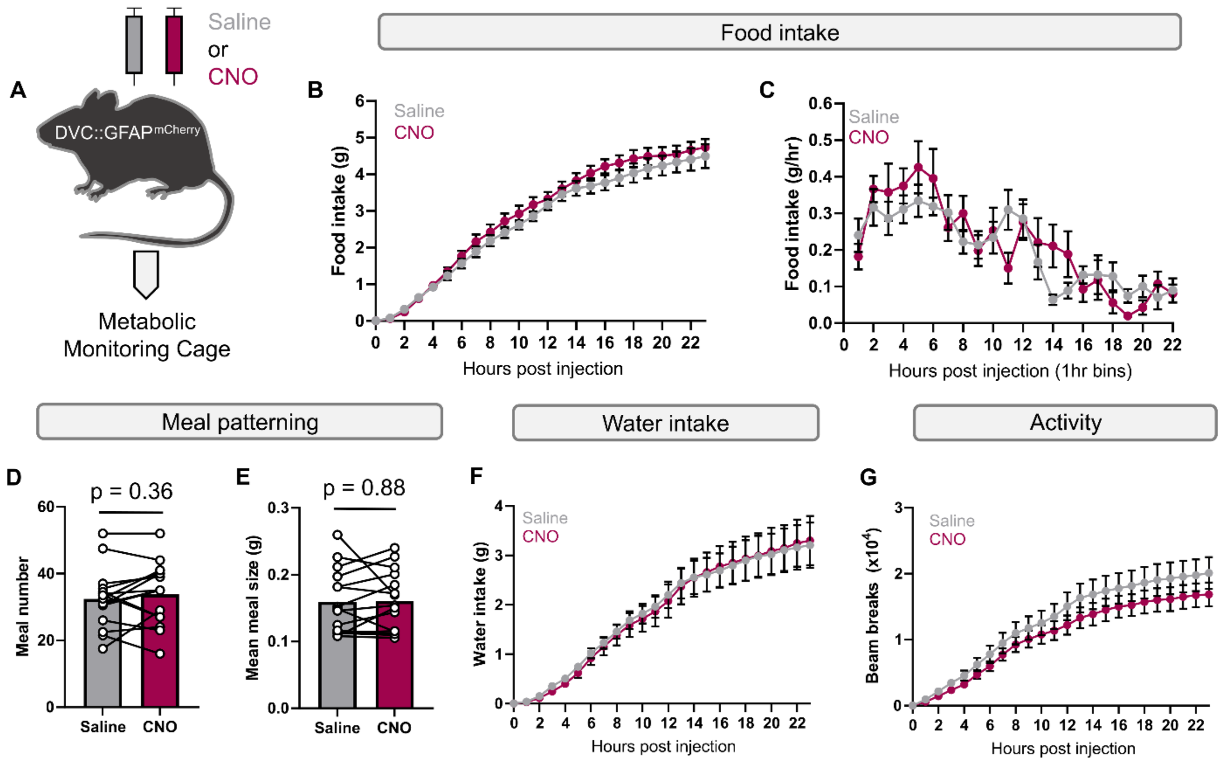


**Supplementary figure 3 | CNO alone does not alter food intake, meal patterning, water intake but has small effect on activity. A,** Schematic of experimental protocol (n=15 mice, 8 female, 7 male). **B,** Cumulative food intake of DVC::GFAP$^{mCherry}$ mice following injection with saline (grey) or CNO (magenta) (two-way RM ANOVA with Sidak's post-hoc test; $p_{treatment}$= 0.24 $F_{(1, 14)}$= 1.51, $p_{time}$< 0.0001 $F_{(23, 322)}$= 249.4, $p_{interaction}$=0.12 $F_{(23, 322)}$= 0.12). **C,** Food intake rate of DVC::GFAP$^{mCherry}$ mice following injection with saline (grey) or CNO (magenta) (two-way RM ANOVA with Sidak's post-hoc test; $p_{treatment}$= 0.33 $F_{(1, 14)}$= 1.01, $p_{time}$< 0.0001 $F_{(22, 286)}$= 11.36, $p_{interaction}$=0.054 $F_{(22, 286)}$= 1.56). **D,** Meal number of DVC::GFAP$^{mCherry}$ mice following injection with saline (grey) or CNO (magenta) (paired t-test). **E,** Meal size of DVC::GFAP$^{hM3Dq}$ mice following injection with saline (grey) or CNO (blue) (paired t-test). **F,** Cumulative water intake of DVC::GFAP$^{hM3Dq}$ mice following injection with saline (grey) or CNO (blue) (two-way RM ANOVA with Sidak's post-hoc test; $p_{treatment}$= 0.88 $F_{(1, 14)}$= 0.022, $p_{time}$< 0.0001 $F_{(23, 322)}$= 45.00, $p_{interaction}$=0.99 $F_{(23, 322)}$= 0.42). **G,** Cumulative activity of DVC::GFAP$^{hM3Dq}$ mice following injection with saline (grey) or CNO (blue) (two-way RM ANOVA with Sidak's post-hoc test; $p_{treatment}$= 0.008 $F_{(1, 14)}$= 9.54, $p_{time}$< 0.0001 $F_{(23, 322)}$= 73.19, $p_{interaction}$<0.0001 $F_{(23, 322)}$= 5.47).

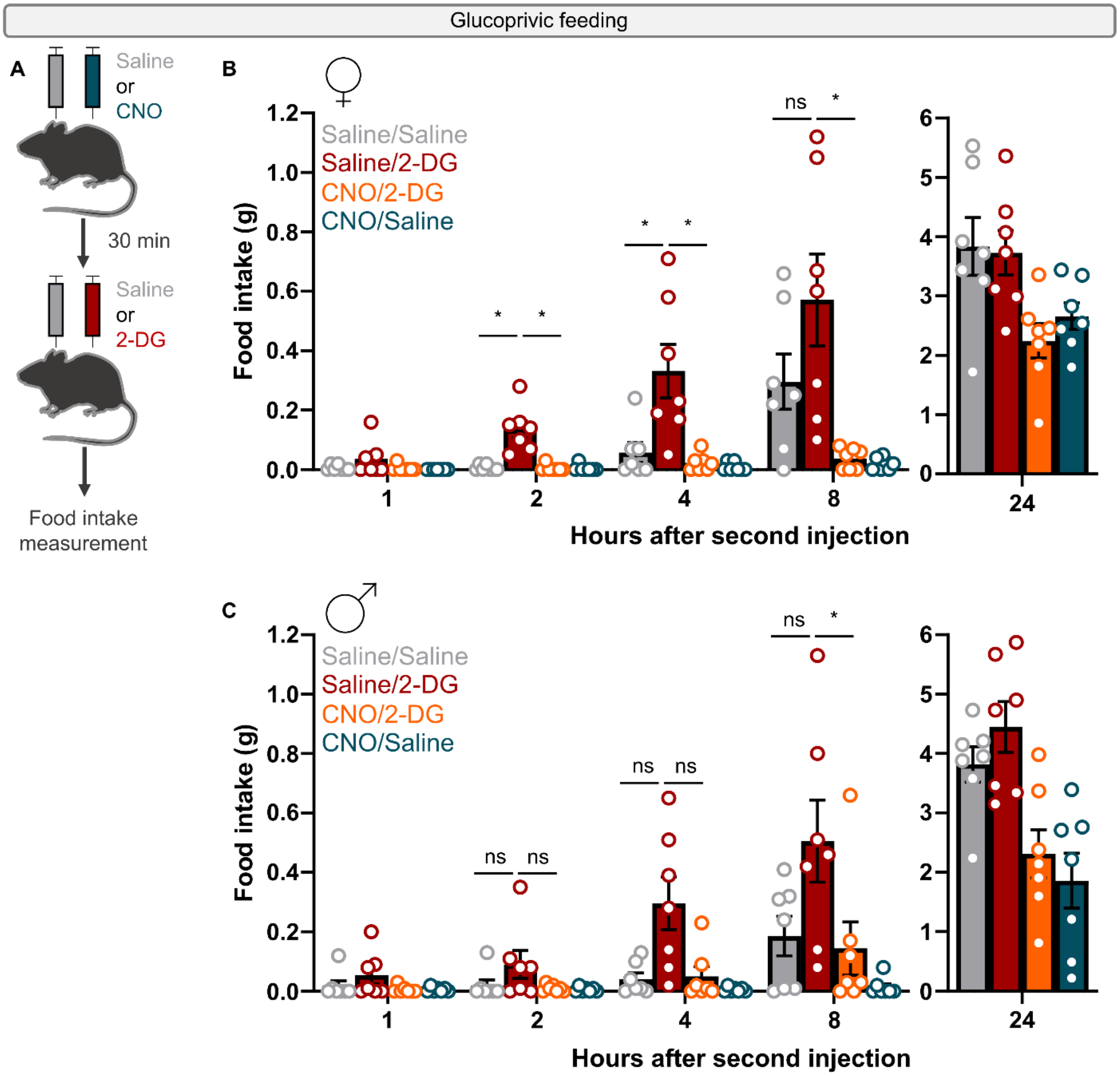


**Supplementary Figure 4 | Glucoprivic feeding stratified by sex. A,** Schematic of experimental protocol. **B,** Cumulative food intake of female mice injected with Saline or CNO followed by saline or 2-DG (n= 7 female mice, two-way RM ANOVA with Geisser-Greenhouse correction, Sidak's post-hoc test; $p_{treatment}$= 0.002 $F_{(2.001, 12)}$= 10.73, $p_{time}$< 0.0001 $F_{(1.095, 6.57)}$= 152.0, $p_{interaction}$= 0.012 $F_{(2.118, 12.71)}$= 6.262). **C,** Cumulative food intake of male mice injected with Saline or CNO followed by saline or 2-DG (n= 7 male mice, two-way RM ANOVA with Geisser-Greenhouse correction, Sidak's post-hoc test; $p_{treatment}$< 0.0001 $F_{(2.363, 13.95)}$= 23.74, $p_{time}$< 0.0001 $F_{(1.067, 6.40)}$= 97.83, $p_{interaction}$= 0.0012 $F_{(1.76, 10.58)}$= 14.49).

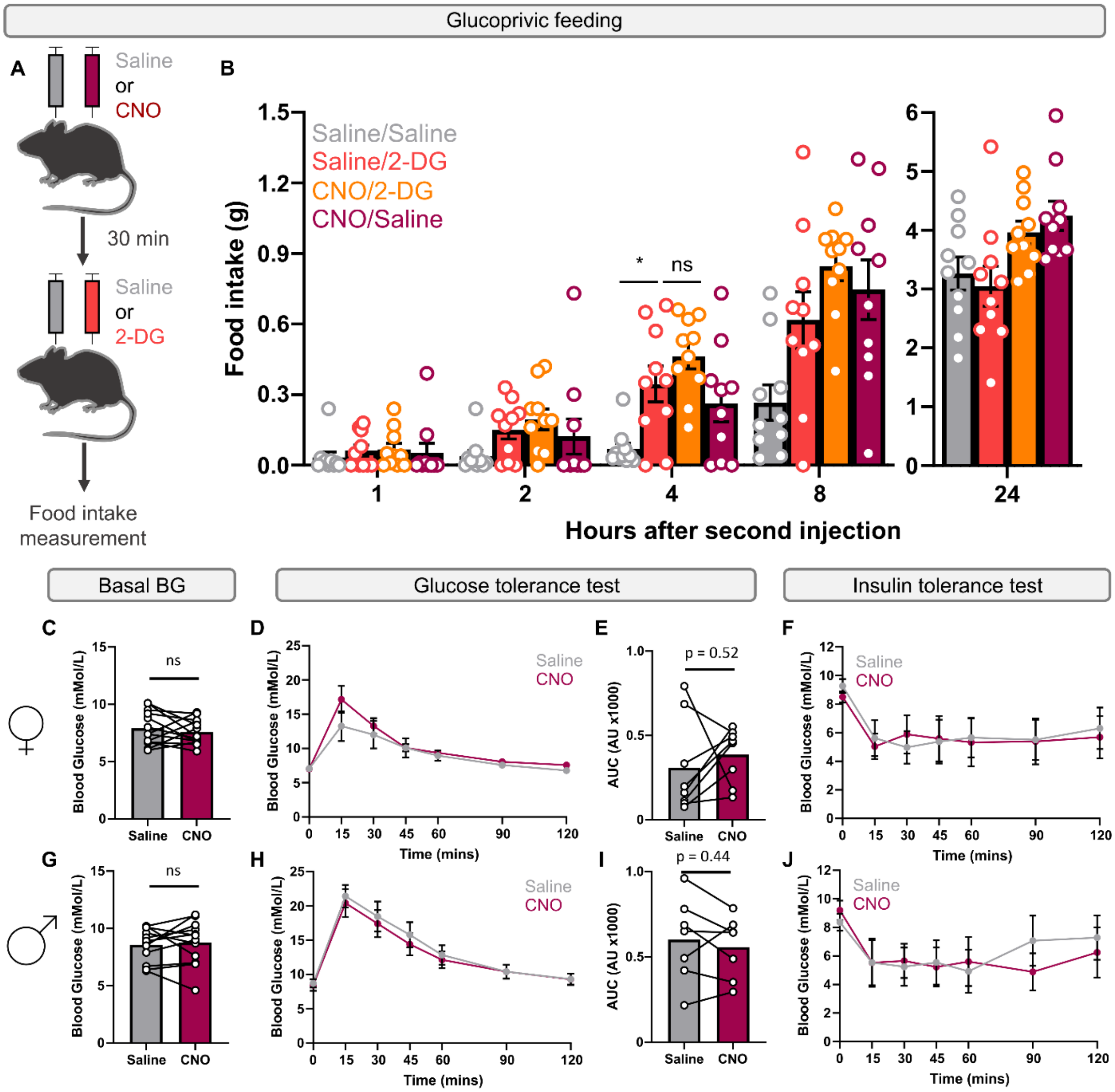


**Supplementary Figure 5 | CNO alone does not influence glucoprivic feeding, basal blood glucose, glucose tolerance or insulin tolerance. A,** Schematic of experimental protocol. **B,** Cumulative food intake of mice injected with Saline or CNO followed by saline or 2-DG (n= 10 mice, 5 female, 5 male, two-way RM ANOVA with Geisser-Greenhouse correction, Sidak's post-hoc test $p_{treatment}$= 0.009 $F_{(2.58, 22.92)}$= 5.19, $p_{time}$< 0.0001 $F_{(1.12, 10.07)}$= 380.3, $p_{interaction}$= 0.006 $F_{(2.52, 22.68)}$= 5.80). **C,** Basal blood glucose (BG) measured 30 minutes after injection with saline or CNO (n = 8 female mice, paired t-test with Bonferroni correction). **D,** Glucose tolerance curve of mice injected with saline or CNO (n= 8 female mice, two-way RM ANOVA with Sidak's post-hoc test; $p_{treatment}$= 0.42 $F_{(1, 7)}$= 0.73, $p_{time}$< 0.0001 $F_{(6, 42)}$= 24.18, $p_{interaction}$= 0.2 $F_{(6, 42)}$= 1.5). **E,** Baseline subtracted area under the curve (AUC) for glucose tolerance test in mice injected with saline or CNO (n= 8 female mice, paired t-test). **F,** Insulin tolerance curve of mice injected with saline or CNO (n= 5 female mice, two-way ANOVA with Sidak's post-hoc test; $p_{treatment}$= 0.83 $F_{(1, 4)}$= 0.05, $p_{time}$< 0.0001 $F_{(6, 24)}$= 4.46, $p_{interaction}$= 0.87 $F_{(6, 24)}$= 0.4). **G,** Basal blood glucose (BG) measured 30 minutes after injection with saline or CNO (n = 7 male mice, paired t-test with Bonferroni correction). **H,** Glucose tolerance curve of mice injected with saline or CNO (n= 7 male mice, two-way RM ANOVA with Sidak's post-hoc test; $p_{treatment}$= 0.44 $F_{(1, 6)}$= 0.68, $p_{time}$< 0.0001 $F_{(6, 36)}$= 54.99, $p_{interaction}$= 0.74 $F_{(6, 36)}$= 0.74). **I,** Baseline subtracted area under the curve (AUC) for glucose tolerance test in mice injected with saline or CNO (n= 7 male mice, paired t-test). **J,** Insulin tolerance curve of mice injected with saline or CNO (n= 5 male mice, two-way ANOVA with Sidak's post-hoc test; $p_{treatment}$= 0.89 $F_{(1, 4)}$= 0.02, $p_{time}$= 0.0001 $F_{(6, 24)}$= 7.48, $p_{interaction}$= 0.36 $F_{(6, 24)}$= 1.16).
